# Hemispheric specialization of functions are tuned by conduction velocities of neuronal propagation in large-scale brain networks

**DOI:** 10.1101/2024.05.02.592033

**Authors:** Neeraj Kumar, Philippe Albouy, Dipanjan Roy, Arpan Banerjee

**Affiliations:** Cognitive Brain Dynamics Laboratory, National Brain Research Centre, NH8, Manesar, Gurgaon 122052, India; CERVO Brain Research Centre, Laval University, Québec, Canada-G1J 2G3; School of AIDE, Center for Brain Science and Applications, Indian Institute of Technology (IIT), Jodhpur 342030, India

**Keywords:** Neural delay, Hemispheric specialization, Auditory networks, Whole-brain model

## Abstract

Analogous to the notion of pleiotropy where one gene can influence two or three unrelated traits, shared structural connectome encompassing both hemispheres of human brain gets functionally segregated along left vs right hemispheres during speech and melody processing, respectively. Here, the underlying neural mechanism is explored with a combined empirical and computational approach wherein, context-specific causal outflow of information emerging from primary auditory cortices (PAC) is shown to be guiding the hemispheric specialization of speech and melody processing while human volunteers listened to naturalistic music while their electroencephalogram (EEG) were recorded. Using a detailed whole-brain connectome model guided by diffusion MRI, individual-specific functional lateralization were predicted at cortical sources representing an extended auditory processing network. High levels of accuracy in prediction can be achieved after optimizing conduction velocity - controller of transmission delays of neural information processing. Thus, a tuning-by-delay mechanism is presented, which when triggered by the spectro-temporal complexity of the task context, guides the geometry of lateralization.

## Introduction

Hemispheric specialization of functions has been established as a fundamentally important empirical phenomenon in neuroscience with substantial clinical relevance (Broca, 1865; Sperry, 1974; Geschwind & Galaburda, 1985; Hellige, 1993; Zatorre & Belin, 2001; Toga & Thompson, 2003; Hecht, 2010; Gotts et al., 2013; Kong, et al., 2018; Brederroo et al., 2019; Karolis et al., 2019). While significance of hemispheric specialization from the perspective of efficient information processing is unanimously agreed on (Corbalis 2017; Güntürkün et al., 2020; for auditory domain see Hickok & Poeppel, 2007; Zatorre, 2022), many studies have identified the lack of mechanistic insights in understanding of how functional specialization occurs at the first place (Gotts et al., 2013; Güntürkün & Ströckens 2017; Brederroo et al., 2019; Karolis et al., 2019; Güntürkün et al., 2020). From a theoretical perspective, the conceptual paradigms of *statistical* and *causal complementarity*, and *input asymmetry* have been used to explain the findings such as increased left hemispheric dominance of language processing and right hemispheric dominance of face processing and motor processing (Brederroo et al., 2019; Gotts et al., 2013). Statistical complementarity suggests that lateralization is an outcome of independent statistical sampling (Liu et al., 2009), causal complementarity and input asymmetry, on the other hand, propose lateralization of one function affects the other. For instance, more left lateralized for language processing is indicative of more right lateralized face processing (Gerrits et al., 2019). The input asymmetry can integrate these frameworks by arguing co-lateralization can emerge selectively because of a hierarchical consolidation of ipsilateral processing from low level to high level cortical structures (Woodhead et al., 2011). Thus, higher activity in left occipitotemporal cortex for higher spatial frequency sinusoidal gratings compared to lower frequencies which have a right ward bias, will lead to leftward bias of processing speech-word stimuli which carry fast spectro-temporal modulations of sound waves.

Earlier studies have indicated that such asymmetry in lateralization schemes can be brought forth by unequal involvement of inter-hemispheric communication for different functions via integrative mechanisms from the vantage point of brain network mechanisms (Gotts et al., 2013, Kumar et al., 2023). Concurrently, observations of specialization in other species having smaller and distinctly different kind of brain topology raises the question whether specialization can solely be explained on the basis of inter-hemispheric transfer, and argue that functional specialization must occur from a set of complex multi-scale processes (Güntürkün et al., 2020). One possible candidate beyond inter-hemispheric transfer may be conduction speeds across the entire connectome that shapes network dynamics which in turn has to encode the spectro-temporal complexity of environmental stimuli (Karolis et al., 2013). Conduction speeds govern the transmission delays in neural communication among brain network nodes and have been identified as a key variable that facilitates the entry and exit to synchronized oscillatory states of the brain (Jirsa & Ding, 2004; Ghosh et al., 2008; Petkoski & Jirsa 2019; Pariz et al., 2021). While, several studies have argued that the changes in transmission delays are of structural origin specifically introduced via myelin modulations along axons (Ghosh et al., 2008; Petkoski & Jirsa 2019; Pathak et al., 2022), empirical evidence points to the existence of functionally tuned synaptic delays (Stange-Marten et al., 2017; Byczkowicz et al., 2019). Functional requirements are set up by a complex system of interactions between internally generated network states and external inputs such as arousal (Vinck et al., 2015), or by neuromodulators (Byczkowicz et al., 2019). Here, a - tuning by delay hypothesis is proposed to conceptualize that while the myelin can provide the basic infrastructure of communications such as expressways to go at high speeds, the functional requirement set by internal awareness system and external stimuli context decides the speed on such paths. Earlier studies have proposed synaptic scaling weights can be indirectly measured from diffusion weighted imaging (dWI) data (Schirner, et al., 2015) and can guide the whole-brain synchrony captured in fMRI and MEG signals (Cabral et al., 2011; Pathak et al., 2022). Hence, it becomes imperative to test that given an individual’s brain structural information, how the task-specific input asymmetries such as speech or melody contexts can be mapped to the conduction velocities with which communication is orchestrated among the auditory network nodes.

Concomitantly, converging evidence suggests that the left A4 (anterior to PAC) is specifically attuned to rapid temporal changes associated with speech processing, while the right A4 is dominant in processing spectral changes associated with melody processing (Albouy et al., 2020; Zatorre & Belin, 2001). Therefore, the lateralization observed in higher-order cortical areas could likely to originate, at least in part from the PAC. Understanding how this oscillatory information is transmitted to higher-order regions is crucial for comprehensive mapping of the complex dynamics underlying auditory perception and consequently to hemispheric specialization. To address this issue, the current article sought to measure the causal flows of information processing using Granger-Gewke causality (Geweke 1982), from source reconstructed EEG time series on human participants when they listened to *a capella* songs and selectively attended to either the speech or melodic content in similar in-principle settings of a previous fMRI study (Albouy et al., 2020). The task involved either discriminating between speech content (same or different lyrics) of two stimuli while melody was varied in delayed-match-to-sample format or discriminating between melody content when speech content was varied. Lateralization indices (LI) were computed for the inflow to each source node after applying Granger-Geweke causality measures on source-level EEG data. Subsequently, to unravel the mechanistic basis of empirical LI patterns, model-predicted LI (referred henceforth as predicted LI) was computed from whole-brain connectome model informed by realistic structural parameters for each individual. The model involved the Kuramoto oscillator framework (Kuramoto 1984) to capture the phase dynamics of an individual source node (Cabral et al 2011; Petkoski et al., 2018; Pathak et al., 2022) coupled through connection matrix generated from diffusion weighted imaging (dWI) data. Conceptually, transmission delays can be set at the gatekeeper node PAC, receiver of auditory sub-cortical output from medial geniculate nucleus (Rauschecker and Tian 2000). The present study also uses a model inversion technique that computes the delays by optimizing a broader parameter space of neural response frequency and synaptic coupling - both within hemispheric and inter-hemispheric, paving the way for individualized prediction of LI required to establish the robustness of the analysis (Seghier & Price, 2018) (see Methods). The frequency-specific tuning of transmission delays was validated by additional auditory steady state (ASSR) recordings an additional control task, where brain networks are entrained sharply to input frequencies of sounds.

## Results

### Behavioral responses

Trials were categorized into speech and melody trials based on the context of discrimination of “speech” or “melody” in consecutive presentations (delayed-match-to-sample) of *a cappella* songs to human volunteers when their electroencephalogram (EEG) data were collected. The percentage accuracy for each cue category was calculated on a participant-wise basis. Participants’ accuracy across both speech (median 97.91%) and melody conditions (median 80.20%) were always consistently above 50% (chance level) (Fig. 1A-II). The difference in accuracy between speech and melody conditions were evaluated using a Wilcoxon signed rank test and was found to be significant at 95% confidence levels (Z=4.71, signedrank =435, p<0.0001).

**Figure 1:**
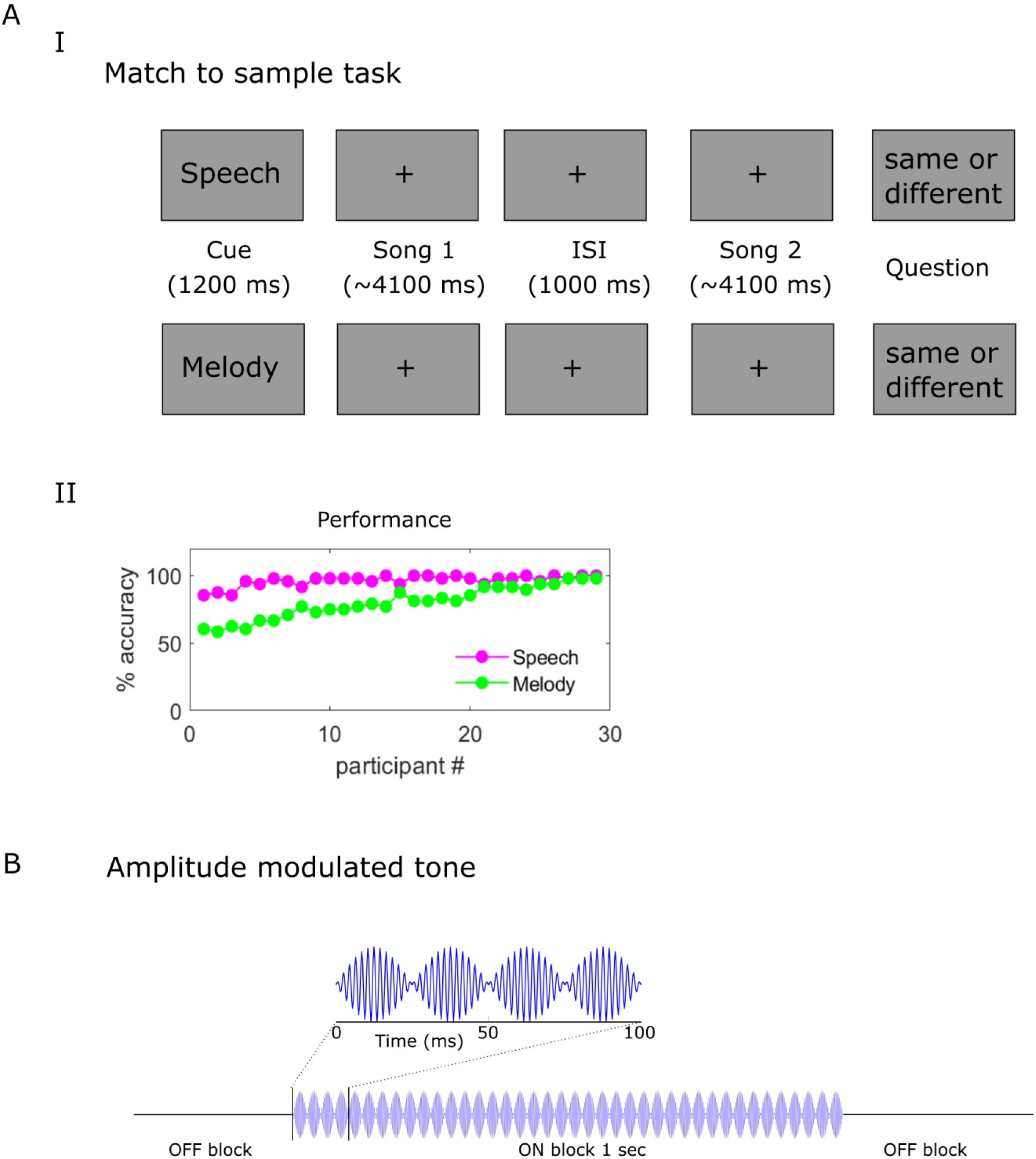
Match to sample task and behavioral responses: (A) (Upper panel) Each trial included a cue, pair of song presentations, and a decision phase, requiring domain-specific (speech or melody) comparisons while ignoring other domain. Participants selectively focused on either speech or melody of a cappella song. (Lower panel) Participant-wise accuracy to correctly discriminate pair of a cappella song based on provided cue. (B) 500 Hz sinusoidal tone amplitude modulated at 40 Hz to generate auditory steady-state responses (ASSR).

### Analysis of EEG signals: Source localization and modelling

The analysis of EEG signals involved a two-pronged approach as outlined in Fig. 2: empirical and theoretical. Empirical EEG data was source localized using their individual head model created from segmented MRI images of their own 3T brain images (MPRAGE). Significant sources were identified via statistical thresholding and and source time series were reconstructed for three auditory conditions, speech, melody and 40 Hz ASSR using eLORETA. Laterality indices (LI) were calculated on frequency-specific net Granger causality outflow from bilateral primary auditory cortices which were estimated for each task condition. Second, theoretical whole brain connectome model was created for an individual’s brain by extracting their diffusion weighted connectivity maps. Numerical simulations were run, by conceptualizing each brain parcel as a Kuramoto oscillator and simulated EEG spectra was generated. The entrainment propagated throughout the entire brain constructed via coupling brain areas (using Desikan-Kiliany parcellation) with strengths derived from diffusion weighted imaging derived structural connectivity (see Methods, as well as Schirner, et al 2015; Pathak et al 2022). Model parameters - a range of realistic conduction velocities that influences the effective time-scales of processing information (scales inversely with time delays in the oscillator model, *τ*), optimal signal frequencies and global synaptic coupling values (k) among the nodes, were predicted using maximum likelihood-based fitting of empirical LIs using model generated LIs. Parameters estimated from this model inversion step were subsequently used for inferring about the underlying neural mechanisms of functional hemispheric lateralization. LI on simulated data was computed by applying net Granger causality on simulated EEG source level time-series using the whole brain connectome model (Fig. 2).

**Figure 2:**
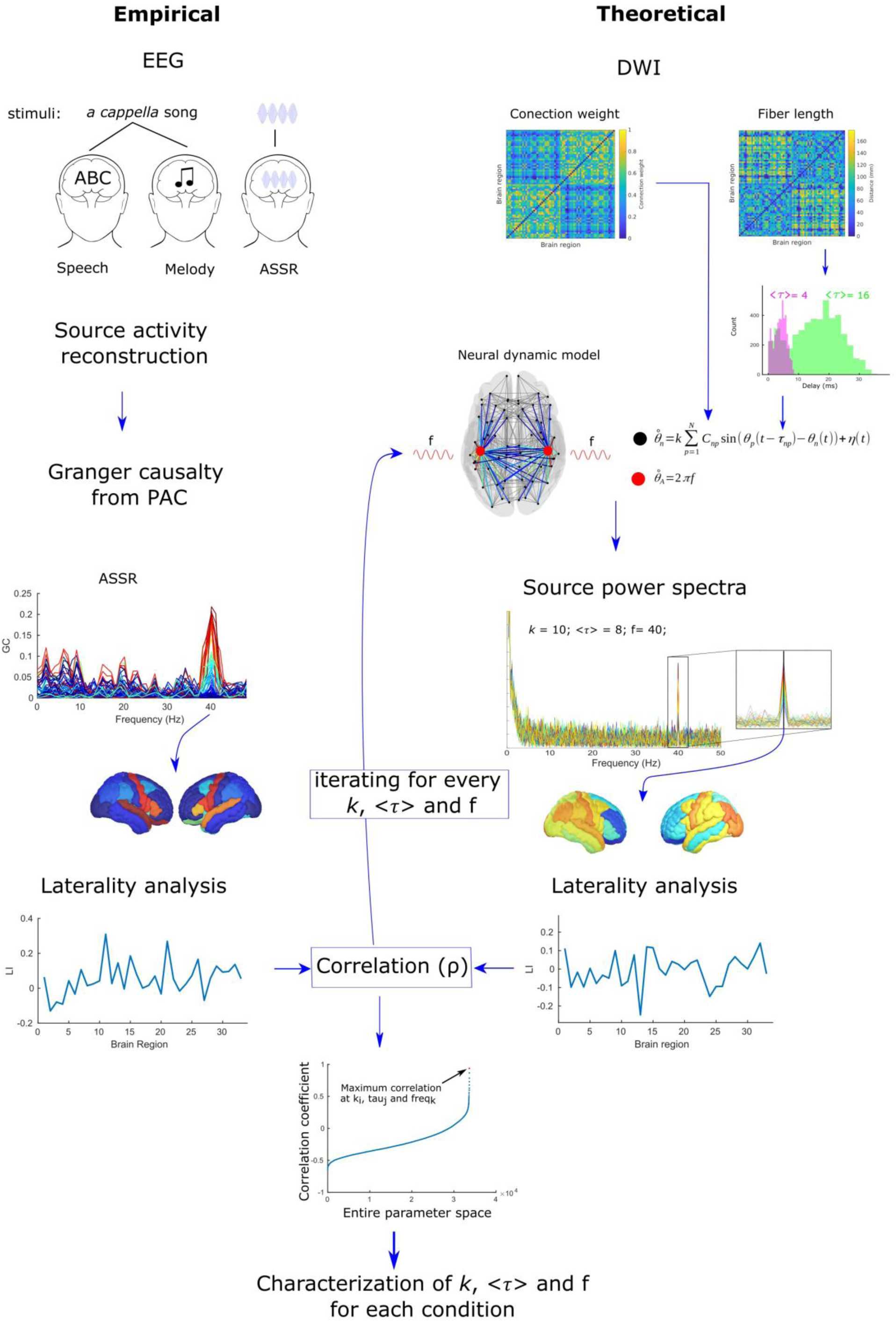
Overview of the methodology: The figure illustrates the pipeline of study, incorporating both empirical and theoretical analyses. The left panel depicts the empirical analysis, highlighting the experimental setup and data processing steps involved in recording neural activity using electroencephalography (EEG) while participants selectively attended to speech or melody stimuli. For visualization purpose, after source localization only ASSR analysis was represented. The right panel represents the theoretical analysis, showcasing the averaged structural connectivity (SC) network that was derived from diffusion weighted imaging (dWI) data. This network served as a constraint for a neural dynamic model to simulate the frequency-specific outflow from the primary auditory cortices from which lateralization indices (LI) can be calculated for specific brain areas.

### Causal outflow of information from primary auditory cortices maps auditory input asymmetries

Cross-spectral matrices at source level were reconstructed using eLORETA at parcels identified by Desikan Kiliany atlas for each participant during speech, melody, ASSR and resting conditions. Multivariate Granger causality analysis revealed significant (p<0.01) causal outflow from bilateral primary auditory cortices to other parcels at distinct frequency ranges: delta, theta, beta and gamma during both speech and melody processing (Fig. 3). The spatially averaged causal outflow across all significantly active sources were similar in the alpha frequency for speech and melody perception but differed across delta, beta and gamma (Fig. 3; Right panel). During the ASSR condition, the significant outflow from bilateral PAC were observed specifically at 40 Hz, whereas speech and melody processing exhibited distinct frequency bands with some sources showing responses at multiple frequency bands (see Fig. 3; left panel and Table 1 for details). Overall, the sources included superior temporal, middle temporal lobe, precentral, Broca’s areas (pars opercularis and pars triangularis), superior frontal, supramarginal and insula. Source in the lateral orbitofrontal and parahippocampal cortex were exclusive to ASSR. Notably, the left pars opercularis showed significant activation across all frequency bands during speech conditions, indicating its involvement in left hemisphere dominance during language processing. On the other hand, the right pars triangularis exhibited significant activation across all frequency bands specifically during melody conditions, suggesting its role in right hemispheric dominance of melody processing (Fig. 3).

**Figure 3:**
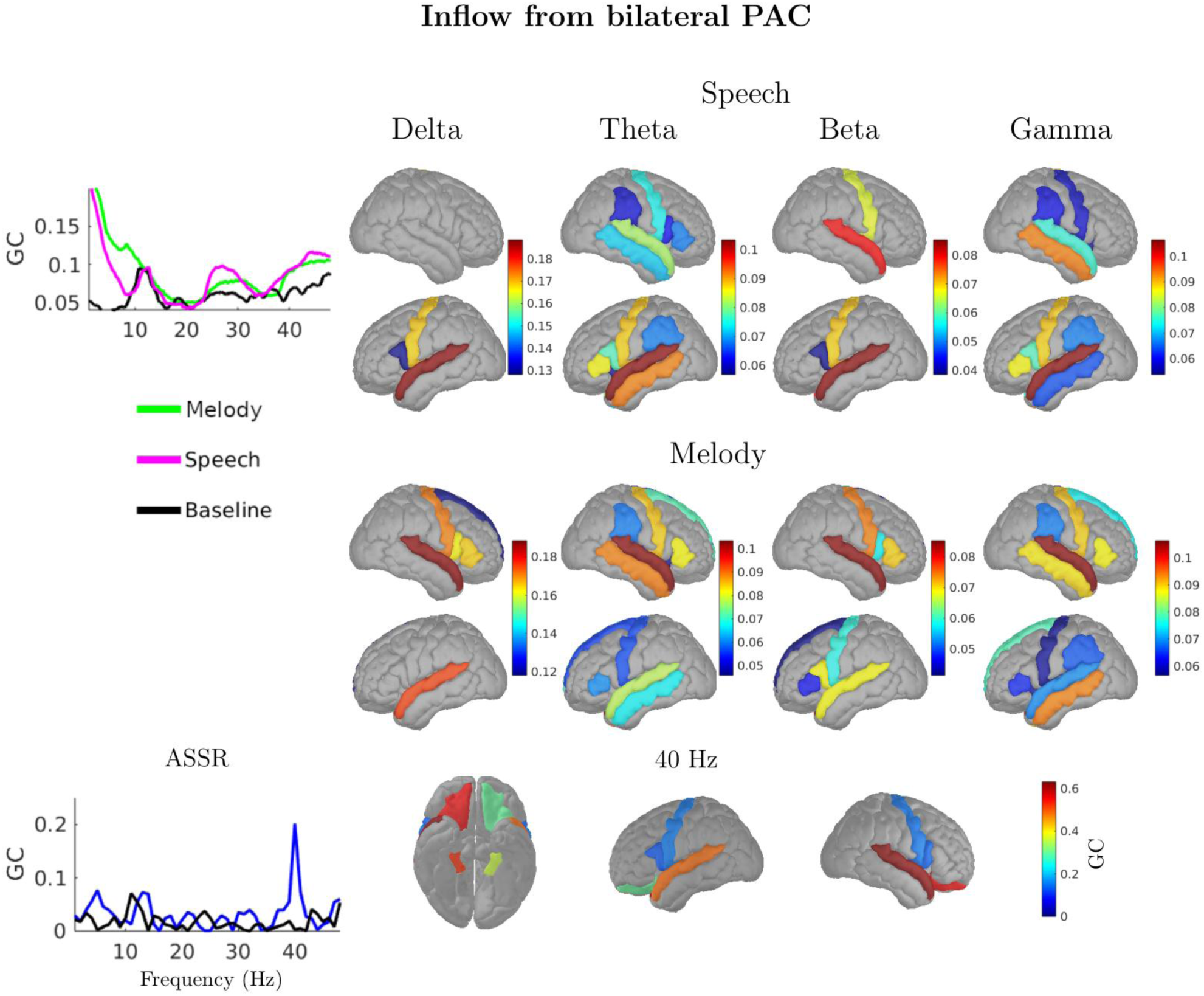
Frequency-specific outflow and target regions of bilateral PAC. The left panel displays the averaged Granger causality spectra, representing outflow from the bilateral primary auditory cortices (PAC) to the target regions. The right panel presents the activation maps of the target regions, averaged for the delta, theta, beta, and gamma frequency bands. For the ASSR condition, only the 40 Hz frequency is represented, reflecting the frequency-specific neural response observed at this frequency.

**Table 1:** Frequency-specific target regions of bilateral primary auditory cortices during different auditory conditions.

| Condition | Frequency Band | Regions |
| --- | --- | --- |
| Speech | Delta | Left Pars Opercularis, Left Precentral, Left Superior temporal |
|  | Theta | Left Middle temporal lobe, Left Pars Opercularis, Left Pars Triangularis, Left Precentral, Left Superior temporal, Left Supramarginal, Left Insula, Right Middle temporal lobe, Right Pars Opercularis, Right Pars Triangularis, Right Precentral, Right Superior temporal |
|  | Beta | Left Pars Opercularis, Left Precentral, Left Superior temporal, Right Precentral, Right Superior temporal |
|  | Gamma | Left Middle temporal lobe, Left Pars Opercularis, Left Pars Triangularis, Left Precentral, Left Superior temporal, Left Supramarginal, Right Middle temporal lobe, Right Precentral, Right Superior temporal, Right Supramarginal, Right Insula |
| Melody | Delta | Left Superior temporal, Right Pars Opercularis, Right Pars Triangularis, Right Precentral, Right Superior frontal, Right Superior temporal |
|  | Theta | Left Middle temporal lobe, Left Pars Triangularis, Left Precentral, Left Superior frontal, Left Superior temporal, Right Middle temporal lobe, Right Pars Triangularis, Right precentral, Right Superior frontal, Right Superior temporal, Right Supramarginal, Right Insula |
|  | Beta | Left Pars Opercularis, Left Pars Triangularis, Left Precentral, Left Superior frontal, Left Superior temporal, Right Pars Opercularis, Right Pars Triangularis, Right Precentral, Right Superior temporal |
|  | Gamma | Left Middle temporal lobe, Left Pars Triangularis, Left Precentral, Left Superior frontal, Left Superior temporal, Left Supramarginal, Left Insula, Right Middle temporal lobe, Right Pars Triangularis, Right Precentral, Right Superior frontal, Right Superior temporal |
| ASSR | 40 Hz | Left Pars Opercularis, Left Precentral, Right Lateral orbitofrontal cortex, Right Parahippocampal, Right Pars Opercularis, Right Precentral, Right Superior temporal |

Lateralization of outflows from bilateral PAC to the other significantly active parcels for different auditory conditions are captured by computing laterality indices for each parcel. The comprehensive list of regions, along with their corresponding 95% confidence intervals and mean LI are provided in Table 2. Certain regions exhibited significant lateralization to either side; where both the upper and lower confidence intervals lie completely on either side of zero (Table 2). Particularly, during ASSR the right hemispheric dominance was present in superior temporal gyrus (STG) with mean LI = 0.13 and lower and upper confidence interval (CI) at 95% significance - [0.08, 0.18]; lateral orbitofrontal (LI = 0.31, CI = [0.17, 0.44]) and parahippocampal (LI = 0.19, CI = [0.05, 0.32]). Analogously, for the speech condition, the left hemispheric dominance in beta band power was present in STG (LI = -0.03, CI = [-0.04, -0.02]), precentral (PrC) (LI = -0.03, CI = [-0.05, -0.01]), pars opercularis (LI = -0.07, CI = [-0.10, -0.05]) and in gamma band power for STG (LI = -0.13, CI = [-0.15, -0.10]), PrC (LI = -0.16, CI = [-0.24, -0.08]), pars opercularis (LI = -0.22, CI = [-0.31, -0.13]). For the melody condition, right hemispheric dominance of beta band was present in STG (LI = 0.10, CI = [0.06, 0.14]), PrC (LI = 0.11, 95% CI = [0.03, 0.18]), pars triangularis (LI = 0.20, CI = [0.05, 0.35]) and at gamma band, in STG (LI = 0.17, CI = [0.12, 0.22]), PrC (LI = 0.17, CI = [0.04, 0.31]), pars triangularis (LI = 0.12, CI = [0.03, 0.22]).

**Table 2:** Confidence Intervals (CI) at 95% significance and mean laterality indices during speech, melody, and ASSR Conditions.

|  | ASSR |  |  | Speech beta |  |  | Speech gamma |  |  | Melody beta |  |  | Melody gamma |  |  |
| --- | --- | --- | --- | --- | --- | --- | --- | --- | --- | --- | --- | --- | --- | --- | --- |
|  | Lo CI | Up CI | Mean | Lo CI | Up CI | Mean | Lo CI | Up CI | Mean | Lo CI | Up CI | Mean | Lo CI | Up CI | Mean |
| Superior temporal | 0.08 | 0.18 | 0.13 | 0.04 | 0.02 | -0.03 | 0.15 | 0.10 | -0.13 | 0.06 | 0.14 | 0.10 | 0.12 | 0.22 | 0.17 |
| Precentral | 0.18 | 0.14 | -0.02 | 0.05 | 0.01 | -0.03 | 0.24 | 0.08 | -0.16 | 0.03 | 0.18 | 0.11 | 0.04 | 0.31 | 0.17 |
| Pars triangularis | 0.13 | 0.27 | 0.07 |  |  |  | 0.40 | 0.03 | -0.19 | 0.05 | 0.35 | 0.20 | 0.03 | 0.22 | 0.12 |
| Pars opercularis | 0.16 | 0.16 | 0.00 | 0.10 | 0.05 | -0.07 | 0.31 | 0.13 | -0.22 | 0.20 | 0.04 | -0.08 |  |  |  |
| Middle temporal |  |  |  |  |  |  | 0.01 | 0.29 | 0.14 |  |  |  | 0.14 | 0.09 | -0.02 |
| Superior frontal |  |  |  |  |  |  |  |  |  | 0.19 | 0.12 | -0.04 | 0.15 | 0.11 | -0.02 |
| Supramarginal |  |  |  |  |  |  | 0.13 | 0.05 | -0.04 |  |  |  | 0.11 | 0.13 | 0.01 |
| Insula |  |  |  |  |  |  | 0.12 | 0.12 | 0.00 |  |  |  | 0.18 | 0.03 | -0.07 |
| Lateral orbitofrontal | 0.17 | 0.44 | 0.31 |  |  |  |  |  |  |  |  |  |  |  |  |
| Parahippocampal | 0.05 | 0.32 | 0.19 |  |  |  |  |  |  |  |  |  |  |  |  |

### Estimation of transmission delays, synaptic scaling and neurally mapped input frequencies

Kuramoto phase oscillator network (Kuramoto, 1984) coupled by heterogeneous coupling derived from diffusion weighted images were employed to model neural oscillations observed in the whole-brain connectome (see Methods for details). Since the goal of the present article is only in the causal outflow from PAC, the input node was accordingly restricted to PAC for the simulations (see Methods for details, equations 3 and 4). Subsequently, the steady state oscillatory responses in each node of this network stems from the inflow of communication from PAC. Hence, the laterality indices (LI) can be computed using the power spectral density at each parcel. By maximizing the correlations between the simulated and empirical LIs, a model inversion strategy was undertaken to estimate transmission delays (*τ*_*opt*_), global coupling (*k*_*opt*_ and neurally mapped input frequencies (*f*_*opt*_). Subsequently, the following issues were addressed. First, the auditory steady state responses were considered, tuned sharply at a specific entraining input frequency to validate if the network model can predict the LIs, tuned at the known frequency, while, readjusting transmission delays and global coupling (Fig. 4). Second, after establishing the validity of the auditory network model, the complex multifrequency EEG response was investigated for speech and melody processing to estimate the transmission delays and global coupling while ensuring frequency selectivity in band specific (delta, theta, beta and gamma) responses and maximizing the LIs (Fig. 4). The estimated parameters can then be used to understand the mechanisms of processing speech and melody by a commonly shared structural auditory network.

**Figure 4:**
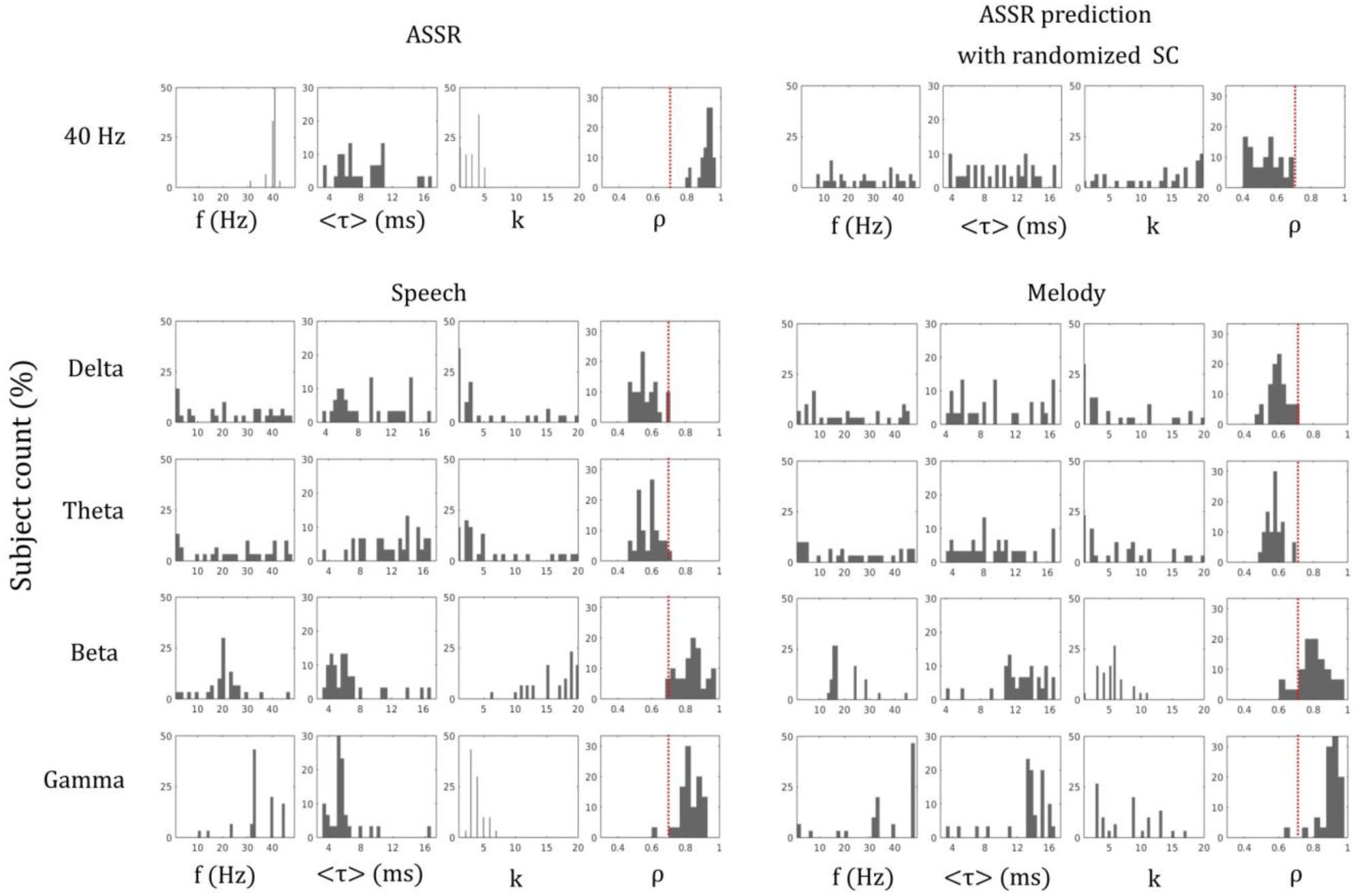
Prediction accuracy of frequency simulation (f (Hz)), global coupling (k), and global delay (*τ* >): Distributions of prediction accuracy of free parameters ASSR, ASSR with shuffled SC, speech, and melody conditions. The x-axis represents the different simulation parameters, while the y-axis represents the prediction accuracy. The first column in each condition demonstrates the effectiveness of the frequency simulation in accurately predicting the neural responses. The last panel shows the correlation coefficients of respective predictions.

### Validation of auditory network model using ASSR

Since, ASSR involves a sharp frequency tuned response, the proposed auditory network model parameters when fitted with a constraining factor of maximizing LI, should be able to detect the frequency of tuned neural oscillations around 40 Hz. Using a realistic parameter search space, *f*_*opt*_ ∈ [1,48], *k*_*opt*_ ∈ [1,20], *τ*_*opt*_ ∈ [3.417], frequency-selectivity of the simulated auditory network was estimated by first constructing a null distribution with 95% confidence bounds for *f*_*opt*_. The control null distribution was created by shuffling the SC matrices, such that each connection weight is essentially a random rational number between 0 and 1, and using this randomized SC to generate neural time series that was used to fit the empirical 40 Hz ASSR. No frequency-specificity was observed from the resultant synthetic data, *f*_*opt*_ was distributed in the whole *f*_*simrange*_. The maximum correlation ranged between 0.4 to 0.7 across all participants (Fig. 4; upper right panel). In contrast, when using the unshuffled SC matrices, the empirical 40 Hz ASSR showed frequency specificity, with a 95% confidence interval of [39.4, 40.9] Hz and maximum correlation values ranging from 0.79 to 0.96 (both > 0.7), thus, confirming the model’s ability to predict empirical responses. The 95% confidence intervals for *k* and *τ* were [2.5, 3.5] Hz and [1.5, 2.2] Hz respectively. The distributions of both *k*_*opt*_ and *k*_*opt*_ followed a normal distribution, assessed using Lilliefors test with k-stat values of 0.24 (p < 0.001) and 0.16 (p = 0.043) and *τ*, respectively (Fig. 4; upper left panel).

### Mechanisms underlying lateralization of causal outflow in beta and gamma frequencies during speech and melody processing

During speech and melody conditions, no frequency-specificity could be predicted by the auditory network model in the delta and theta frequency bands (Fig. 4; bottom panels). In the delta range, the 95% confidence intervals of *f*_*opt*_ for speech and melody were [17.6, 29.8] Hz and [15.7, 26.9] Hz, respectively. Similarly, in the theta range, the 95% confidence intervals for speech and melody were [19.2, 30.8] Hz and [16.0, 28.1] Hz, respectively. Therefore, the *k*_*opt*_ and *τ*_*opt*_ values for these frequency ranges are not considered reliable. On the contrary, frequency-specificity was observed in beta and gamma frequency bands (see Fig. 4). For speech conditions, the 95% confidence interval of *f*_*opt*_ in beta band was [17.2, 23.7] Hz, and for melody [17.6, 22.8] Hz. In the gamma band, the 95% confidence interval of *f*_*opt*_ was [31.0, 41.7] Hz for speech, and [31.1, 37.4] Hz for melody. These CIs indicated that the model correctly captured the frequency-specific activity of empirical data within the beta and gamma bands. The values of *k*_*opt*_ and *τ*_*opt*_also showed normal distributions for these frequency bands, indicating a smooth and gradual transition in the model’s behavior required for a consistent and reliable model performance. The complete list of 95% confidence intervals for frequency, *k*_*opt*_, and *τ*_*opt*_ as well as the results of the normality tests, are present in Table 3a. The paired t-tests showed the presence of overall differences in neural processing characteristics between speech and melody stimuli (Fig. 6A). The findings revealed a significant difference in the k parameter [t = 3.05, p = 0.003], indicating distinct global coupling between the two conditions. Moreover, there was a significant difference in the tau parameter [t = 11.32, p < 0.0001], suggesting variations in temporal delays during the processing of speech and melody stimuli (Fig. 6A). These analyses shed light on the global neural dynamics and temporal characteristics associated with different auditory stimuli.

**Table 3a:**
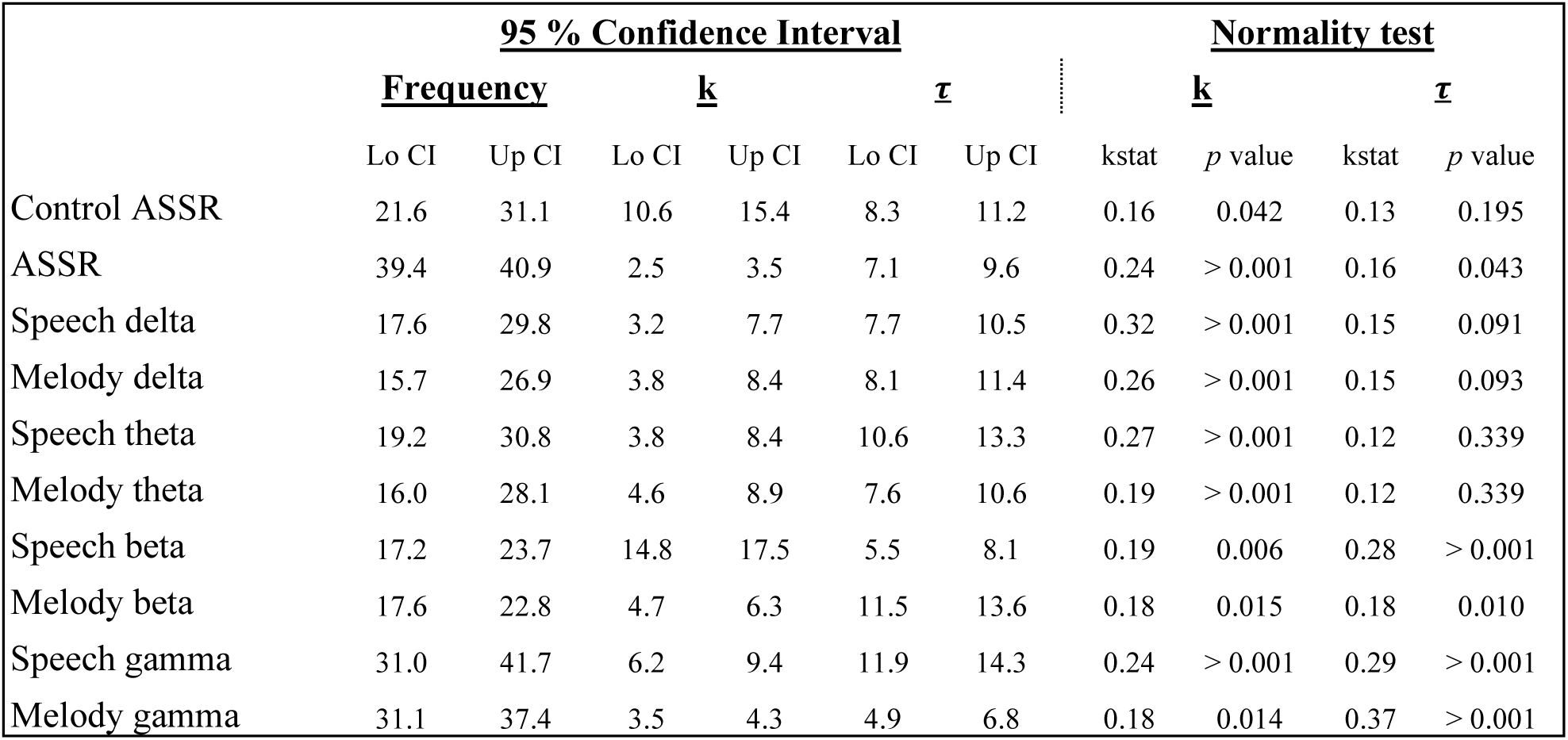
Prediction specificity and normality of free parameters.

### Individual-specific prediction of functional lateralization in auditory processing networks

Linear regression analyses was conducted in the beta and gamma frequency ranges to evaluate the extent of prediction for individual regions found significant in the empirical data. Beta and gamma were the chosen frequency of interest since the correlation between empirical and simulated data were significant for these two bands (Fig. 4). Most regions significantly predicted lateralization in the empirical data, with the exception of the supramarginal gyrus during speech condition in the gamma band. Fig. 5 A show the respective prediction score for the individual brain region. Overall, the prediction scores for the ASSR condition were greater than the speech and melody condition. When both the upper and lower confidence intervals of a region’s laterality index (LI) for both the simulated and empirical data fell on the same side of zero, the model was considered to successfully predict the side of lateralization for that region. Consistent with the empirical conditions, several regions exhibited lateralization. Particularly, during the ASSR condition, right hemispheric dominance was predicted by the superior temporal gyrus (STG), parahippocampal cortex, and lateral orbitofrontal cortex (Fig. 5B). During speech and melody conditions, the STG predicted left hemispheric dominance during speech and right hemispheric dominance during both beta and gamma frequencies. The precentral gyrus predicted the side of lateralization during speech beta and gamma, and in melody during beta frequencies. Additionally, the pars opercularis predicted left hemispheric dominance during speech gamma conditions.

**Figure 5:**
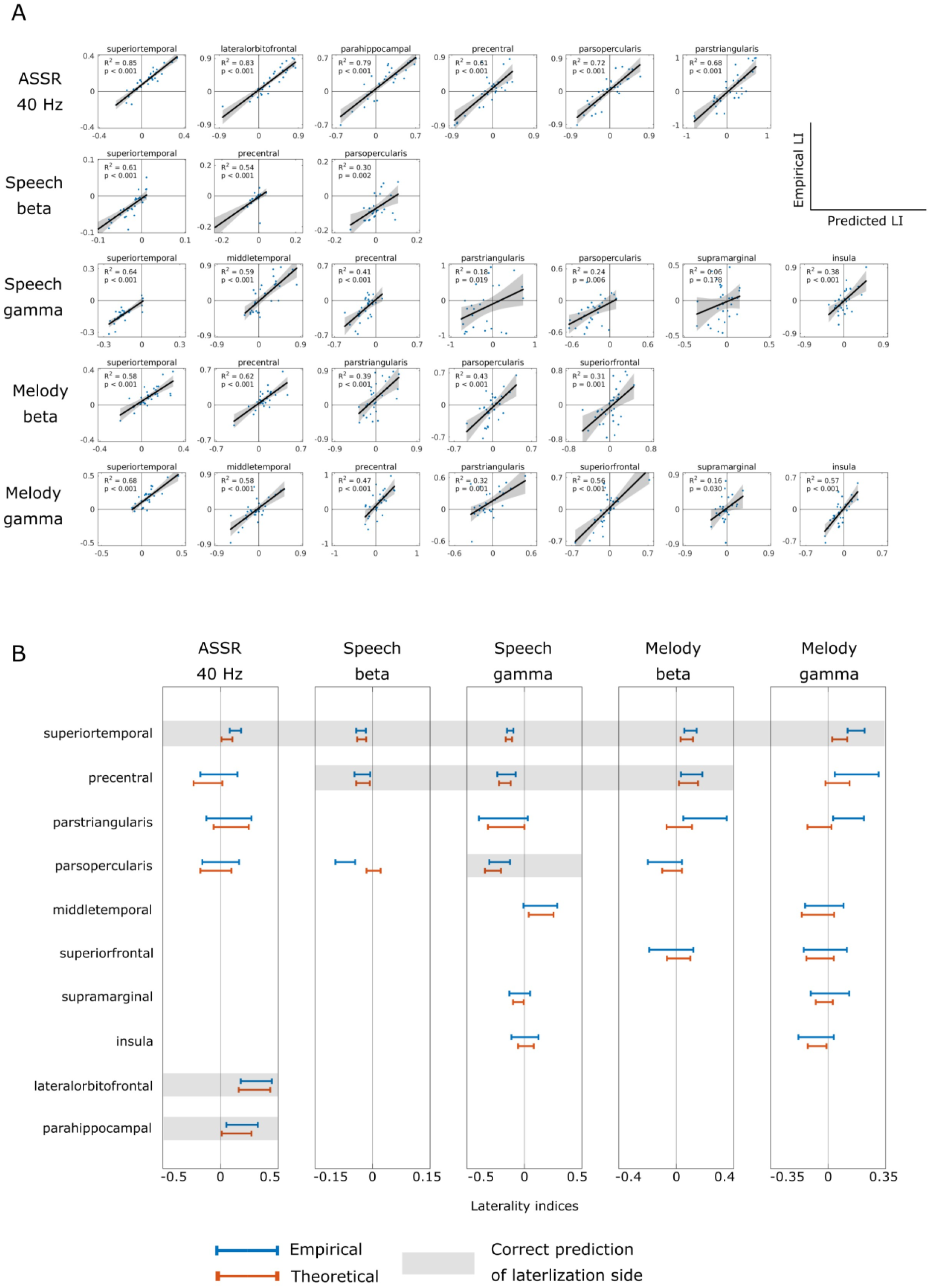
Lateralization prediction accuracy of individual regions: (A) Scatter plot shows the linear fitting of empirical LI (Y-axis) against predicted LI (X-axis). Each dot represents the LI of one participant. The analysis focuses on beta and gamma frequencies for speech and melody conditions, as well as 40 Hz for the ASSR condition. (B) Confidence intervals for LI, the confidence intervals for the LI during empirical (blue) and predicted (orange) analyses. The shaded background indicates cases where both empirical and predicted LIs fall on either side of zero, indicating accurate prediction of the lateralization direction.

Interestingly, a systematic decrease in R-squared values for individual regions was observed along functional hierarchy of information processing. When arranging the regions in order of auditory processing hierarchy, namely superior temporal gyrus (STG), middle temporal gyrus (MTG), insula, precentral gyrus, Broca’s area, and supramarginal gyrus, for both speech and melody conditions, the degree of prediction exhibited a consistent linear decrease in the gamma frequency band (Fig. 6B and Fig. 6C). This pattern was observed in both the speech condition [*r*^2^ = 0.93, p = 0.001] and the melody condition [*r*^2^ = 0.93, p = 0.001]. Note that, similar analysis in the beta frequency band was not feasible due to the limited number of regions. However, in the beta range the highest prediction accuracy was present in STG for speech and PrC for melody condition.

**Figure 6:**
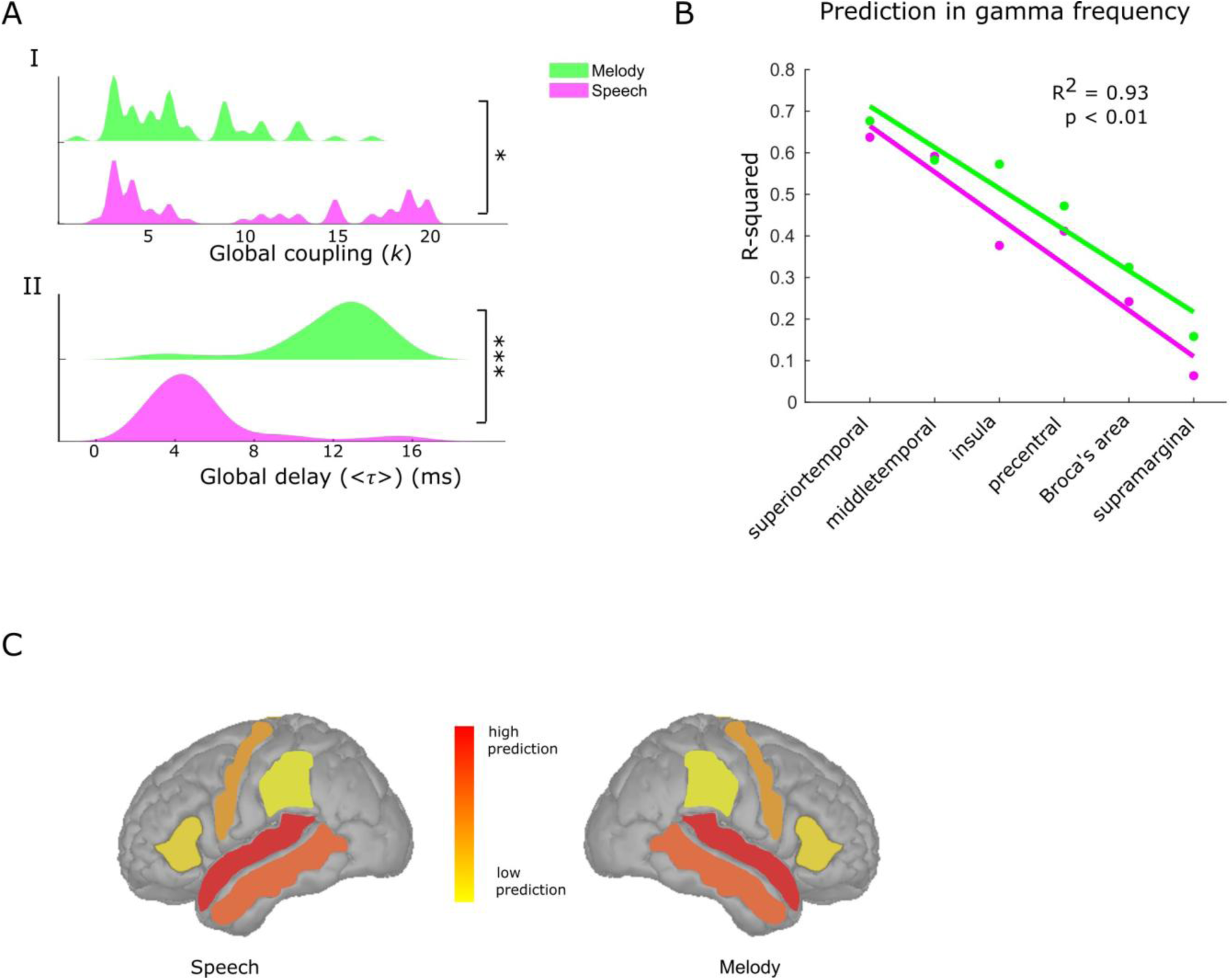
(A) Distribution of global coupling and global delay for beta and gamma frequencies in speech and melody conditions. The distributions are visualized using kernel smoothing with a bandwidth of 0.3 to enhancing the clarity of the patterns. (B) Decrease in prediction accuracy along the auditory hierarchy in gamma range in the speech and melody conditions. (C) Prediction accuracy in homologous auditory areas segregate along sensory processing hierarchies for speech and melody processing.

Finally, whether the extracted parameters of global coupling and delay were correlated with behavioral measure of accuracy in speech and melody contexts were explored. Although accuracy was different across speech and melody conditions and reported in an earlier subsection, none of the model parameters were found significantly correlated with accuracy when Spearman rank correlation was used with p< 0.05 after correcting for multiple comparisons using False Discovery Rate - Benjamini-Hochberg (Table 3b).

**Table 3b:** Spearman’s rank correlation coefficients (corrected) between accuracy in speech/ melody contexts with global parameters (k, *τ*).

| Accuracy |  | Speech |  |  | Melody |  |
| --- | --- | --- | --- | --- | --- | --- |
|  |  | *r* | *p* |  | *r* | *p* |
| Beta | **k** | -0.38 | 0.36 |  | -0.04 | 0.82 |
| | $\tau$ | -0.18 | 0.83 | | 0.26 | 0.80 |
| Gamma | **k** | -0.06 | 0.83 |  | 0.09 | 1.00 |
| | $\tau$ | -0.21 | 0.84 | | 0.08 | 0.94 |

## Discussion

The present study uses a combination of empirical EEG recordings and biophysically detailed whole-brain connectome modelling approach to propose that hemispheric lateralization of complex cognitive functions can emerge from modulation of the conduction speeds during task-specific network processing. Such time-delay tuned neurodynamics in a distributed auditory processing network leads to the empirical observation of hemispheric lateralization for speech and melody processing - introduced as task contexts via a psychophysical paradigm design. A common frequency-specific outflow of information communication from PAC is reported for both speech and melody conditions, reaching to distinct lateralized regions reflecting the domain-specific organization of the auditory system. This enhances the well-established hemispheric dominance of left hemisphere for speech, right hemispheric bias for melody, elucidating the important role of PAC guided causal flow in hemispheric specializations. To further understand the origins of this flow patterns and subsequent lateralization, a structurally-guided frequency-specific neural model was conceptualized to predict an individual’s lateralization index (LI) from their brain responses. The novel approach of whole-brain connectome modeling to explain task induced electrophysiological response during speech, melody, and ASSR conditions led to a prediction of LI of an individual participant in homologous brain areas at a high level of accuracy. Thus, the second notable aspect of current study is the individual level parametrization of the whole-brain connectome model to capture variability in brain structure across humans. By integrating individual diffusion weighted imaging (dWI) data into the neural model, participant-specific variability in structural constraints is considered to explain the inter-participant variability of neural dynamics (Seghier & Price, 2018). The frequency-specificity of the model was validated and characterized for the neurophysiologically relevant conduction velocity and global coupling characteristics of each auditory condition ASSR, speech and melody. Thirdly, from a systematic investigation of the model predicted parameter space of conduction speeds, a dissociation of network transmission delays for speech and melody processing was established, stemming from the input asymmetry in spectro-temporal complexity of these two task contexts. This supports the existence of a tuning-by-delay mechanism in cortical networks for routing contextual information from identical stimuli into two hemispherically distinct homologous areas. In fact, such tuning-by-delay mechanism can explain not only the conundrum of lateralization of speech and melody but also in other domains such as auditory-motor coordination where right hand (controlled by the left hemisphere) seems more agile to fast finger tappings to auditory beat compared to left hand (controlled primarily by right hemisphere) which can follow slower frequency tappings more accurately (Pflug et al., 2019). Finally, even though there was a specific focus on the role of the primary auditory cortices (PAC) and their outflow in governing hemispheric specialization, the modelling tool itself can be tweaked to predict other functional and effective connectivity measures as well and hence have tremendous application scenarios. In following sections, these issues are discussed in more detail.

### Hierarchical and domain-specific organization of the auditory system

Several researchers have established that the auditory connectome is hierarchically organized (Kaas & Hackett, 2000; Rauschecker & Tian, 2000) with the PAC being the first cortical structure to receive ascending auditory inputs exhibiting a rich repertoire of oscillatory activity (Gourévitch et al., 2020; Hackett, 2011). Albouy and colleagues demonstrated the sensitivity of the left A4 to speech features and the right A4 to melody features (Albouy et al., 2020). On the other hand, there is substantial evidence of left hemispheric specialization for speech and right hemispheric specialization for music (Zatorre & Belin, 2001; Zatorre et al., 2002). In the present study, the laterality profile of inflow in higher-order auditory regions from bilateral PAC complies with well-established hemispheric dominance of left hemisphere for speech, right hemisphere for melody and ASSR (Fig. 3 and Fig. 5B), suggesting observed hemispheric specializations could be attributed to the feed-forward flow from PAC. This contributes to the understanding of the hierarchical organization of auditory processing, as it reveals the flow of oscillatory information from lower-level sensory regions to higher-order cognitive regions. The oscillatory activity in PAC and the neural markers of its coordination with higher order areas seems to be specific to the nature of the auditory stimuli (Fig. 3). These specific communication dynamics across different regions are required for binding of disparate features, the segmentation of auditory streams, and the formation of hierarchical representations (Kösem & van Wassenhove, 2017; Plakke & Romanski, 2014).

Although the oscillatory patterns observed during speech and melody conditions are similar, characterized by frequency-specific outflows from the bilateral PAC, the specific regions involved in receiving these oscillations differ between speech and melody conditions. This suggests that the observed oscillatory patterns do not encode generic processing of acoustic signal dynamics but instead capture specific linguistic or musical features. The present experimental paradigm ensures selective attention of participants to the speech or melody aspects of ecologically valid auditory stimuli (*a cappella* songs), thereby eliminating confounding effects of signal properties. The involvement of distinct regions in processing these acoustic environments indicates specialized and domain-specific organization within the auditory system (Angulo-Perkins & Concha, 2019; Peretz et al., 2015; Scott & McGettigan, 2013; Xie et al., 2018). Furthermore, the presence of homologous areas in the opposite hemisphere, such as the superior temporal gyrus (STG), motor cortex, and Broca’s area, suggests a bilateral representation and processing of auditory stimuli. Homologous areas share an evolutionary history and tend to perform similar functions, necessitating communication to coordinate their respective functions (Agcaoglu et al., 2018; Karolis et al., 2019; Wan et al., 2022). This parallel processing mechanism enables the brain to handle the complementary nature of speech and melody, two most important auditory communication skills. By distributing the cognitive load, the brain enables simultaneous processing when speech and melody are presented together, as in the case of ecologically valid *a cappella* songs (Angulo-Perkins & Concha, 2019). The identified respondent regions of the primary auditory cortices (PACs) in current study during both speech and melody conditions are consistent with previous literature and widely recognized as auditory-related regions (Giraud & Poeppel, 2012; Hickok & Poeppel, 2007; Morillon et al., 2010). The regions in the left hemisphere are known for processing various linguistic aspects, including phonological analysis, syntactic structure, semantic processing, and articulation. Conversely, the right hemisphere is implicated in processing non-linguistic aspects of auditory stimuli, such as melodic and tonal structures, pitch variations, temporal patterns, and rhythmic patterns in music (Zatorre & Belin, 2001; Zatorre & Gandour, 2008). Additionally, the ability of a single region to receive input at multiple frequencies highlights the brain’s capacity to decode and integrate distinct aspects of oscillatory activity for coherent representation of auditory percept. This multiscale hierarchy integrates motor, syntactic, and working memory information, contributing to the construction of a comprehensive auditory scene. Thus, functional integration at these cognitive levels can be identified using EEG activity with high temporal resolution, that opens up the path for studying many functions in detail.

### Role of structural constraints in lateralized network dynamics

Structural symmetry of the brain, particularly the patterns of anatomical connectivity between brain regions, plays a crucial role in shaping the processing of information in brain networks (Honey et al., 2007; Sporns, 2010). In the auditory domain, auditory functions of speech and melody is postulated to be hinged in structural connectome (Mišić et al., 2018). In order to dig deeper into the mechanistic principles of such functional asymmetries, the prediction of LI using brain network topology from real human brain were compared to that of LI predicted from a network where brain connectivity is organized randomly, but frequencies, delay and coupling can be tuned (Fig. 4). The neural model incorporating participant specific structural connectivity (SC) while tuning for delay, brain response frequencies and couplings achieved the highest prediction accuracy in the 40 Hz ASSR condition (Fig. 4). The high model accuracy sheds light to the fact that brain responses can be entrained to an oscillatory response mode that captures the collective dynamics of large-scale network via modulation of time delays (functional parameter) for a given structural connectivity (participant specific parameter). The high degree of predictability of LIs in homologous brain areas during ASSR response (ρ = [0.79 0.96]) provide candidates that the model inversion techniques will also be capable of detecting the response frequencies, coupling and transmission delays in situations where spectro-temporal complexity increases. Thus, all in all, the usage of biologically realistic network topology in the neurodynamic model expanded the capability of the models (Sanz Leon et al., 2013; Schirner, et al 2015) and also underlines the importance of structural connectivity shaping mode dynamics of auditory processing. Furthermore, high prediction in motor regions in beta frequencies could be attributed to the “hard-wiring” of auditory–motor interaction as reported earlier (Morillon et al., 2010) and further consolidates the role of structural asymmetries in shaping up function. The linear decrease in prediction accuracy across hierarchical brain regions (Fig. 6B) further support the general view that auditory processing areas follows a posterior to anterior hierarchy from simple to complex processing from primary sensory to complex cognitive areas. Although some recent advances in whole-brain modelling suggests geometry is also an important factor (Pang et al., 2023), however, such is currently out of scope of the present model but certainly remains a candidate for future explorations. Finally, the synaptic scaling used by brain regions can have structural origins owing to fibre densities among regions and functional origins measured via functional and effective connectivity. In presented model, while the connection weights (*C*_*ij*_) are estimated from fibre densities global coupling is a parameter that uniformly scales all functional associations such as in high and low attention scenarios, thus regulating synchronization strength. By modulating these factors, the brain can finely adjust the degree of coordination between neural signals across different areas.

### Tuning by delay hypothesis for hemispheric lateralization

An important point to note is that brain structure is static in an observational window of sessions and days while functions are not. While basic features of auditory processing can be explained solely based on the network properties of structural connectivity, more abstract and higher-level features of the environment, such as meaning, context, and emotion require a more complex and dynamic interplay between different regions of the brain (Bauer et al., 2020; Berger et al., 2019; Di & Biswal, 2019). These findings hint that the higher-level processing or other factors beyond phase-based interactions contribute to propagation of delta and theta oscillations (Pandey et al., 2022). On the other hand, the processing of meaning in language requires the coordination of neural activity between regions involved in phonetic, lexical, and semantic processing, as well as regions involved in attention, working memory, and executive control (Kazanina & Tavano, 2023). Similarly, the processing of emotion in music requires the coordination of neural activity between regions involved in auditory processing, pitch perception, and emotional valence (Gnanateja et al., 2022; Wang et al., 2023; Yurgil et al., 2020). Although, the distinct functional requirements are supported by the structural connectivity between these regions, nonetheless, brain accomplishes such higher cognitive processing by precisely manipulating the timing and coordination of neural signals through two key factors: conduction velocities and synaptic scaling (Pathak et al., 2022; Petkoski & Jirsa 2019). Conduction velocities, which controls the transmission delay of neural impulses travel along nerve fibers, play a pivotal role in governing the temporal aspects of neural communication. (Ghosh et al., 2008; Pflug et al., 2019; Pathak et al., 2022). Varied conduction velocities facilitate the emergence of distinct time delays, thereby shaping the arrival and integration of neural information across disparate regions (Cariani & Baker, 2022; Pariz et al., 2021). This temporal coding ensures the precise synchronization necessary for intricate processing tasks (Ibrahim et al., 2021; Petkoski et al., 2018). Hence, despite the underlying static structural connectivity within the brain that changes over the time-scale of months or years, modulation by conduction velocities and coupling strength enables the generation of a diverse range of time delays and patterns of information propagation.

The computational model revealed that differences in conduction velocities contribute to the functional lateralization observed in speech and melody processing (Fig. 6A). The faster conduction velocities in the left hemisphere, specialized for speech, enable the precise analysis of temporal changes associated with speech processing, while the slower conduction velocities meaning greater time window allows to capture fine spectral resolution of melody in the right hemisphere. These dichotomies could be attributed to enhanced myelination thereby enhanced transmission speed in the left hemisphere than right hemisphere (Seldon, 1981, 1982) which has also been reported for motor coordination (Pflug et al., 2019). Most importantly observed differences in conduction velocities between speech and melody stimuli in present study align with previous findings that speech exhibits rapid temporal changes and music shows fast spectral changes (Zatorre & Belin, 2001). The role of dichotomous or distinct categories within a system could be the causes of development of separation of cognitive processes or functional specialization observed in dichotomies. Moreover, as suggested earlier, brain’s parallel processing mechanism allows it to effectively handle the complementary aspects of speech and melody, which are two crucial forms of auditory communication. By distributing the cognitive load, the brain enables simultaneous processing when speech and melody are combined, as seen in ecologically valid a cappella songs. Somewhat subtle, yet an important point to distinguish is while myelination may set in place the overall network tuning frequencies within a hemisphere, the functional input asymmetry decides the spatial geometry (left vs right) of lateralization.

One interesting observation which highlights the limitation of the present approach is that the extracted functional variables of conduction speeds and global coupling were not able to predict the inter-subject variability in accuracy although the model was set up constraining the LIs (Table 3b). This indicates the different level of LIs used for constraining the model across participants may not have a task specific advantage at least from the perspective of accuracy. However, since this was not one of the fundamental goals of the study, a more nuanced analysis can be undertaken by conducting a detailed study varying auditory segment lengths, difficulty index of discrimination, etc on the extracted neurofunctional parameters in future. An alternative way to interpret may be that the parameters extracted are purely of sensory origin without any task benefits. Another limitation of the present Kuramoto framework approach to study the tuning-by-delay hypothesis is that the role of cholinergic signalling (Picciotto, et al., 2012) or other factors such as impact of extracellular current flow (Abodollahi & Prescott, 2024) which can also affect the conduction velocity of spikes along networks remains out of scope. Future studies can be constructed along these lines to identify the more physiologically fine-grained mechanisms which can cause the conduction velocity to change beyond myelin constrained delays of structural origins embedded in networks.

In summary, highlight of this study is mapping the dichotomy or dualistic nature of the auditory system interactions at systems-interaction level. In the case of acoustic signals, viz., *a cappella* songs, the auditory system utilizes two divergent but related (temporal and spectral) cues embedded in a common signal to dissociate perceptual experiences in the form of speech and melody, highlighting the dichotomy of auditory system. The presence of common oscillatory outflow in homologous brain regions is demonstrated and hints at “spatial” dichotomy. The common oscillatory signatures observed in the brain may represent shared underlying neural generators and gives rise to distinct hemispheric dominance. Furthermore, the variations in conduction velocities result in different temporal processing characteristics, enabling the specialized processing of speech and melody in their respective hemispheres highlighting a “temporal” dichotomy. Thus, the findings highlight the interplay between functional dynamics and structural connectivity in shaping auditory processing and importance of dissociation of spatial and temporal scales of processing. Hence, the present study contributes to the understanding of hemispheric specialization and provides an overarching mechanistic explanation of speech, melody, and tonal auditory perception in a common framework. Speech processing involves the encoding of fast temporal changes of auditory stimuli. On the other hand, the melodic processing of musical notes unfolds in slower time-scale to bind temporally far away events. Thus, the perception of speech and melody is determined by the organization of oscillatory activity in the PAC, as well as the subsequent propagation dynamics to downstream auditory networks along the input asymmetries which ultimately leads to hemispheric specialization.

## Methods

### Participants

Thirty healthy participants (14 males, 16 females, age range 22–34 years old; mean ± SD = 27 ± 3.14) participated in this study. All participants were right-handed, as confirmed by the Edinburgh Handedness Questionnaire with a cut-off score of 60-100, and reported no history of audiological, neurological or psychiatric disorders. All had normal or corrected-to-normal visual acuity. Informed consent was obtained from all participants in a format approved by the Institutional Human Ethics Committee (IHEC) of National Brain Research Centre. All participants were fluent in at least two languages, Hindi and English, with some having knowledge of another language of Indian origin.

### Auditory stimuli

Two types of auditory stimuli were used - 1.) *a cappella* songs (Albouy et al., 2020) and 2.) amplitude modulated tones to generate auditory steady-state responses (ASSR). The first set of auditory stimuli consisted of 100 *a cappella* songs prepared by Albouy and colleagues, and have been used and described in more detail in their earlier reports (Albouy et al., 2020). Particularly, the a cappella songs were composed of 10 tones, each with an identical rhythm, and ten English sentences composed of 10 syllables each. Sentences were modified to adjust with the rhythm of the melodies. All melodic and sentence materials were combined in all possible combinations to form 100 a cappella songs. The resulting set of 100 songs had a mean duration of 4.1 seconds. The second set of auditory stimuli were pure sine wave tones of frequency 500 Hz, amplitude modulated at 40 Hz (Fig. 1B). The sound intensity level was maintained at 60 dB SPL for all stimuli.

### Paradigm and experimental sessions

Participants underwent two separate sessions: an MRI session in which T1-weighted structural MRI images and diffusion weighted imaging (dWI) data were acquired, and an EEG session during which electrode locations were also obtained using a 3D digitizer. The EEG session always took place on a different day and was scheduled after the MRI session.

The EEG session consisted of 3 blocks. First, the participants underwent 5 minutes of EEG recording in a resting state while fixating on a cross displayed at the center of the screen. Second, after being verbally instructed about the task participants were binaurally presented with pairs of a cappella songs via earphones and were instructed to compare them. To ensure that participants focused their attention on speech or melody content during the presentation of a cappella song, they were engaged in a match-to-sample task (Fig. 1A: Upper panel). The trial started with a visual cue indicating the relevant domain (sentence or melody) for 1200 milliseconds. Subsequently, the first song was presented, followed by a 1000 millisecond interstimulus interval (ISI), and then the second song was presented. Throughout the entire duration of the songs and ISI, a fixation cross was presented at the center of the screen. Following the presentation of the second song, participants were presented with a question on the screen, requiring them to assess whether the two speech contents were identical or different, irrespective of the melody to which they were sung in ‘speech’ trials. In the case of ‘melody’ trials, this process was reversed, with the judgment based solely on the melody, independent of the sentence content. Participants were required to respond by selecting the right arrow key on the keyboard when the domain was the same and the left arrow key for different. The response window lasted for 2000 milliseconds, and participants received no feedback. The total duration of a trial was approximately 12 seconds, and the total duration of the experiment was 20 minutes. There were four types of combinations included: (1) both speech and melody were different in the song pair, (2) speech was the same but the melody was different, (3) speech was different and the melody was the same, and (4) both speech and melody were the same. Each combination had 24 trials, resulting in a total of 96 trials presented. The sentence/melody questions and same/different responses were pseudo-randomly presented, with each stimulus type uniformly distributed over the entire experiment. Presentation software (Neurobehavioral Systems, Berkeley, CA, USA) was used to control stimulus presentation and record participants’ responses. Continuous electroencephalography (EEG) was recorded in a noise attenuated isolated room using a BrainVision Recorder acquisition system, which included an actiCHamp module with 63 active channels placed according to the International 10-20 electrode placement system. SuperVisc electrolyte gel (EASYCAP) was used to establish contact between the EEG sensors and the scalp. Continuous EEG data were acquired at a sampling rate of 1 kHz, and the impedance of each sensor was kept below 10 kΩ. The reference electrode was positioned at the vertex (Cz), and the forehead (AFz) electrode was selected as the ground. The electrode locations were obtained with respect to three fiducials at the nasion and left and right preauricular points using a 3D digitizer (Polhemus Inc., Colchester, VT, USA).

### EEG Preprocessing

Initial EEG data analysis was performed with EEGLAB (Delorme and Makeig, 2004) for removing artifacts from the data, followed by customized MATLAB scripts (www.mathworks.com) and FieldTrip (http://fieldtriptoolbox.org) toolbox for spectral analysis and source analysis. Firstly, using EEGLAB continuous EEG data from each auditory block and resting state were bandpass filtered between 2-48 Hz and independent component analysis was employed to eliminate eyeblinks, muscular, and heartbeat artifacts. The ASSR block trials were epoched for a duration of 1 second from the onset of stimuli, while for the *a cappella* song block, hit trials were identified and segregated into two groups for speech and melody trials and epoched for a duration 4 seconds from the onset of song. In addition, non-overlapping 1 second and 4 second (50% overlapping) time segments from the resting state data were epoched to be utilized as the baseline condition for the ASSR and song blocks, respectively. After detrending the trials, threshold-based artifact rejection was applied at ± 80 μV on trials. Following the preprocessing steps, an average of 97 trials (out of 100) from the ASSR block, 81 trials (out of 96) from the speech block, and 69 trials (out of 96) from the melody block were retained. The preprocessed EEG data were then re-referenced to a common average reference and downsampled to 250 Hz for further processing.

### MRI-DWI acquisition and analyses

The T1-weighted structural MRI images were acquired on a Philips Achieva 3.0 T MRI scanner with the following parameters: TR = 8.4 ms, FOV = 250 × 230 × 170, flip angle = 8°, 170 sagittal slices, and voxel size of 1 × 1 × 1 mm. The fiducials were marked at the nasion, left, and right pre-auricular points using Vitamin E capsules to be used in EEG sensor coregistration with MRI for source localization. The initial preprocessing of MRI images and volumetric parcellation for individual participants were performed using Freesurfer, based on the Desikan-Kilinay atlas (http://surfer.nmr.mgh.harvard.edu/). Diffusion weighted Imaging (DWI) data was acquired through a single-shot echo planar imaging in 3.0 T Philips Achieva Scanner. The DWI sequence was performed with TR = 8800 ms, TE = 75 ms, FOV = 210 × 210 × 128 mm, flip angle = 9°, matrix size = 104 × 104, slice thickness = 2 mm, no gap, 64 axial slices, and voxel size of 2 × 2 × 2 mm. The diffusion was measured along 32 non-collinear directions using a *b*-value of 1000 s/mm^2^, with a b = 0 image included as the first volume of the acquisition.

### Image processing and structural connectivity

The processing of diffusion data and T1 MRI data to construct structural connectome (SC) was performed in FSL and ANTS based MRtrix (http://mrtrix.org/), BrainSuite (https://brainsuite.org/), and FreeSurfer. Structural connectivity estimation was performed employing the BATMAN pipeline implemented in MRtrix software. Particularly, probabilistic tractography was performed based on the Constrained Spherical Deconvolution (CSD) algorithm (Tournier et al., 2004, Tournier et al., 2007). The CSD algorithm evidently outperforms the diffusion tensor model (DTI) in regions containing crossing fibers, which DTI cannot capture due to its ellipsoid approach to fiber orientation estimation. The initial preprocessing of diffusion-weighted images included denoising to estimate the spatially varying noise maps (Veraart et al., 2016b, Veraart et al., 2016a), unringing to remove Gibb’s ringing artifacts was performed in MRtrix (Kellner et al., 2016) and bias field correction of diffusion images (Tustison et al., 2010). Then MRI images are preprocessed for co-registration with diffusion data. Therefore, firstly, skull-stripping of MR images was performed by the Cortical Surface Extraction Tool (Sandor et al., 1997; Shattuck et al., 2001), followed by correction of gain variation by the Bias Field Corrector (BFC) software (Shattuck et al., 2001), part of the BrainSuite. Particularly, it was used to correct image intensity non-uniformities in magnetic resonance images, which can cause confounding effects in tissue classification. BFC estimates a correction field based on a series of local estimates of the tissue gain variation and corrects for the non-uniformity field by dividing the original image by the estimated tri-cubic B-spline. Thereafter, BrainSuite Diffusion Pipeline (BDP) was used to correct motion and geometric distortions of diffusion images inherent to Echo-Planar Imaging sequences (Bhushan et al., 2012), co-registration of diffusion and anatomical images, and obtaining b-values and b-vectors to be used for further estimation of response functions (Haldar & Leahy 2013; Shattuck et al., 2002). Subsequently, to determine the diffusion orientation within each voxel, a response function was derived using Dhollander’s algorithm from representative tissue types of the brain i.e., white matter, gray matter, and cerebrospinal fluid (Dhollander et al., 2016). Thereafter, Fiber Orientation Distributions (FOD) were estimated using the CSD algorithm. Intensity normalization was performed to correct for global intensity differences, and a whole-brain tractography was generated using probabilistic tractography in tandem with biologically plausible Anatomically Constrained Tractography algorithm to generate 20 million tracts seeded in gray matter (Smith et al., 2012). The tractogram was then coregistered onto individual MRI, and a subset of one hundred thousand tracts was filtered out to reduce CSD-based inherent bias in overestimation of longer tracks by Spherical-deconvolution Informed Filtering of Tracts (SIFT) algorithm (Smith et al., 2013). Finally, the connectome was parcellated by mapping the tracts onto MRI data, defined according to the Desikan Kiliany atlas (Desikan et al., 2006), that was segmented in FreeSurfer (https://surfer.nmr.mgh.harvard.edu/). The resulting 68 x 68 parcellated whole-brain structural connectivity (SC) weights and fiber lengths matrices were representative of the number of streamlines and mean streamline length between each node pair, respectively. Both SC matrices were normalized by the inverse of the two node volumes for scaling the track count. The SC matrix and fiber length matrix were both symmetric matrices and values at the diagonals of structural connectivity and fiber length matrix representing self-connectivity and length with self, respectively, were set at zero.

### Source-level analysis of EEG data

To estimate frequency-specific oscillatory source activity, exact low-resolution brain electromagnetic tomography (eLORETA), a distributed source modeling method that calculates the current source density across the brain volume by minimizing the surface Laplacian component of the spatial filter (Pascual-Marqui, 2007) was used. Notably, eLORETA has the advantage of not requiring any assumptions about the number of underlying sources and is highly effective in controlling false positives during source detection (Halder et al., 2019). Therefore, source analysis was performed using eLORETA implemented from the FieldTrip toolbox (Oostenveld et al., 2011).

For forward modelling, MRI data from each participant was segmented into three tissue types, and meshes were created with 3000, 2000, and 1000 vertices for brain, skull, and scalp, respectively. The boundary element method (BEM) was employed to generate the realistic volume conduction model using the OpenMEEG package (Fuchs et al., 2001; Gramfort et al., 2010). Subsequently, the EEG channels were aligned with the BEM model based on 3 fiducials present in both Polhemus data and MRI. Thereafter, leadfields were created for 3 orthogonal directions using the sensor positions and the volume conduction model. The location of vertices within the leadfield was defined based upon the participant-specific T1 MRI previously parcellated according to Desikan-Killiany atlas in FreeSurfer (Desikan et al., 2006), with an average resolution of 4 mm.

Subsequently, the cross-spectral for each frequency band (2-48 Hz) were computed from the preprocessed sensor level data over the entire time window for each condition for ASSR, speech, melody, and baseline conditions. Frequency-domain spatial filters were computed from combination of the forward model and cross-spectral matrix for each condition. Separate spatial filters were constructed for ASSR, speech, and melody conditions, with 1-second and 4-second rest trials serving as baseline conditions, respectively. To reduce dimensionality, principal component analysis was performed on the spatial filters of vertices belonging to individual parcels, resulting in the spatial filters of 68 regions of interest. The trial-wise sensor-level Fourier-spectra were then projected to the spatial filter, obtaining participant and trial-wise distribution of source level Fourier coefficients across the brain volume for each condition.

### Granger causality analysis

From the source the reconstructed Fourier coefficients of entire frequency range were obtained. Thereafter, participant-wise multivariate Granger causality (GC) was calculated in the spectral domain to determine the causal outflow from PAC to whole-brain (Brovelli et al., 2004; Dhamala et al., 2008; Granger, 1969). To calculate the pairwise Granger causality (GC) from PAC to X regions at frequency (*f*), a nonparametric spectral matrix factorization was performed using Wilson’s algorithm. This factorization of the cross-spectral density allowed for the determination of the transfer function *H* and the noise covariance matrix ∑. Subsequently, the pairwise GC can be expressed as follows:

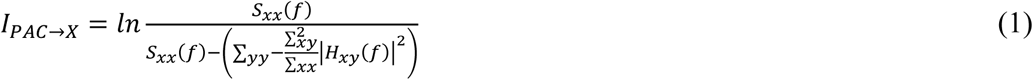

where *S*_*xx*_(*f*) represents the total spectral power (auto-spectrum), while the denominator represents the intrinsic power of the “effect” signal. The intrinsic power is calculated by subtracting the causal contribution from the total power (Geweke, 1982, Dhamala et al 2008).

The GC spectra of speech and melody condition were divided into respective frequency bands, i.e., delta (2-4 Hz), theta (4–8 Hz), alpha (8–13 Hz), beta (13–30 Hz), and gamma (30–48 Hz) bands. Significant frequency bands were identified from non-parametric statistical testing (Brovelli et al., 2004). Particularly, 1000 permuted data sets were generated by independently shuffling trials between resting condition and auditory conditions. Thereafter, GC was computed, and the maximum GC value was selected over the frequency range from each permuted data set (Ding et al., 2006). Subsequently, a null distribution consisting of all GC values was constructed from the shuffled data set. GC bands in unshuffled data were considered statistically significant when the observed GC value reached beyond the 99^th^ quantile value (p = 0.01) of the null distribution. The multiple comparison problem was handled by Bonferroni correction.

### Laterality analysis

To evaluate the extent of asymmetry in outflow from the bilateral primary auditory cortices (PAC) during auditory conditions, the Laterality Indices (LI) were calculated of the inflow to each area (Kumar et al., 2023). The LI for each node of the auditory network was then calculated as the difference between the total GC inflow (from bilateral PAC) to the right region and the left region, divided by the sum of the inflows to both regions.

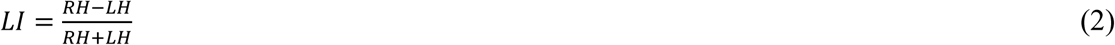

LI values range from +1 to -1, indicating complete right dominance and left dominance, respectively, while a value of 0 suggests a bilaterally symmetrical response. Statistical significance was of the LI distribution across participants were determined by computing 95% confidence intervals.

### Network model of neural activity

Computational modelling of the propagation of oscillatory activity from primary auditory cortices to specialized regions of the brain during speech, melody and ASSR conditions was undertaken using a simplified computational model, retaining the essence of key physiological parameters. The propagation of oscillation can be conceptualized as unidirectional interactions originating in PAC to form a functional neural network. Such networks communicate via phase-based synchronization (Kuramoto, 1984) and were shown to be the processing units of cognition (Cabral et al., 2011; Pathak et al., 2022). Hence, the Kuramoto model was employed to simulate the source time series of each phase oscillator equivalent to phase dynamics of one parcel (Kuramoto, 1984; Cabral et al., 2011; Pathak et al., 2022). Hence, a network of 68 coupled Kuramoto oscillators was simulated, following Desikan-Killiany parcellation scheme, also comprising bilateral primary auditory cortices (PAC). Each other oscillators had intrinsic noise (*η*), which was randomly drawn from a distribution with a mean of zero and a standard deviation of one, reflecting the natural variability of oscillatory dynamics in the brain. The remaining two oscillators situated in the primary auditory cortices and have fixed intrinsic frequencies that correspond to their oscillatory responses represented by red nodes inside glass brain in figure 2 right panel. This framework allowed the unidirectional propagation of entrainment from PAC to the specific nodes and consequent enhancement of spectral power of those nodes at the frequency of PAC stimulation (Fig. 2; Right panel). To make the model bio-physiologically realistic, information about structural connectivity was incorporated in the model by having the coupling between oscillators determined by strength of anatomical connections (fibre thickness in dWI, see section “Image processing and structural connectivity”). The time scale separation for processing of “fast” speech and “slow” melody could be achieved by a propagation time-delay (*τ*) in the model such that the current phase *θ*(*t*) is dependent on its interaction with the past phase *θ*(*t* − *τ*). Inadvertently, *τ* can be conceptualized to be emerging from a joint contribution of the functional processing in the auditory network and the constraints posed by myelination, although the former is the only quantity that can change in a participant and in a session specific way. Thus, the dynamics of each oscillator *θ*_*n*_ are governed by the following equation:

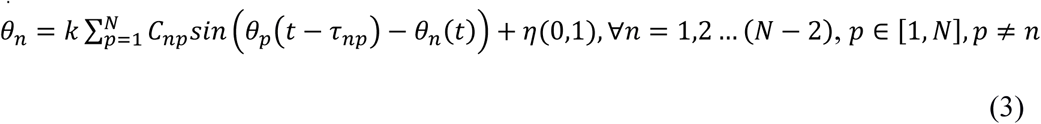

The coupling strength matrix *C*_*np*_ was normalized between 0 and 1 such that the maximum strength among connections was 1 and, *k* is the global mean coupling strength used to scale all the coupling strengths; **τ**_*np*_ represents transmission delay for the propagation of information between two nodes. Thus, 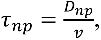 where for a bio-physiologically realistic communication speed (*v*) is 5 − 20 *m*⁄*s* in the adult primate brain (Ghosh et al., 2008), mean *τ* ranged from 3.4 to 17 when *D*_*np*_ was fibre length between nodes *n* and *p*. Therefore, the dynamics of phase (*θ*) at any node is a function of its anatomical strength, distance with other nodes and propagation speed *v*, which is set be the nature of functional processing. While equation (3) describes the dynamics of 66 non-auditory nodes, the phase dynamics of bilateral auditory cortices (auditory nodes) are defined as,

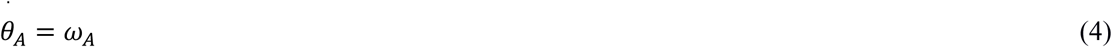

where *ω*_*A*_ = 2*π* ∗ *f*_*A*_, (*f*_*A*_) representing the frequency of auditory nodes.

### Participant-wise model fitting

The participant-by-participant laterality indices obtained from the empirical Granger-causality estimation were fitted with the LI’s computed from analyzing synthetic neural time-series generated by the network model (equations 3, 4). The system of differential equations (equation 3,4) was numerically integrated using the Euler integration method for 250000-time points with a step size (*dt*) of 0.0001 representative of 25 seconds duration. Sine of (*θ*) obtained at each node which represents a simulation of neural time series at EEG source level. Subsequently, the power spectral density were calculated at 68 nodes. For each combination of frequency (*f*), global coupling (*k*), and time delay (*τ*) model simulation, simulated LI were calculated from power spectral density obtained from each 68 parcels (equation 2). An optimization algorithm was run to obtain a set (*f*_*opt*_, *k*_*opt*_, *τ*_*opt*_), at which the Pearson correlation between simulated and empirical LI reaches a global maximum (Fig. S1 A). Hence, a model inversion was achieved for the following ranges of each parameter, *f*_*opt*_ ∈ [1,48]; *k*_*opt*_ ∈ [1,20]; *τ*_*opt*_ ∈ [3.4,17] in steps of 0.4. The frequency window of interest was guided by the observation window of current study, the range of *k*_*opt*_ and *τ*_*opt*_chosen has been shown earlier to represent biophysically relevance (Ghosh et al., 2008; Cabral et al 2011).

To process a specific auditory task, phase-based communication in an oscillatory network can be achieved by adjusting the combination of these free parameters (*f*_*opt*_, *k*_*opt*_, *τ*_*opt*_). While the fitting was constrained by maximizing Pearson correlation between empirical LI and model generated LI, frequency-specificity of these optimal parameters model need to be studied to ensure that the simulated neural responses align with the observed empirical frequency-specific enhancements in Granger causality (GC) outflow from the bilateral primary auditory cortices (PAC) (Fig. S1 A). Hence, the model inversion was carried out independently for the major frequency bands, delta (2-4 Hz), theta (4–8 Hz), alpha (8–13 Hz), beta (13–30 Hz), and gamma (30–48 Hz) for speech and melody and also specifically at 40 Hz for ASSR condition. Importantly, it was required to ensure the frequency specificity of the model i.e., the significant distribution *f*_*opt*_ across participants should fall into the corresponding frequency range. For instance, the confidence interval of *f*_*opt*_ to predict empirical theta lateralization should fall into the theta range and so on. Hence, to establish a valid comparison between empirical EEG frequency and *f*_*opt*_ that can establish the frequency selectivity of the model, correlations between empirical and simulated LI were calculated across a range of probable model frequencies (*f*_*simrange*_) and the 95 % confidence bound of resultant *f*_*opt*_was estimated. Since, the outflow from PAC during speech and melody existed in the broadband frequency range rather than a single frequency unlike the model simulation, making the comparative assessments between model and empirical results were not straightforward resulting to spurious non-unique values of frequencies and other parameters during the model inversion process. Hence, 40 Hz ASSR condition where the signal displays a salient entraining frequency becomes a further validation ground to assess whether the model can be frequency-selective (See Fig. S1 A). Outflow from bilateral PAC was shown earlier to be present only at 40 Hz during 40 Hz ASSR response (Kumar et al., 2023). Hence, for the purpose of current study, demonstration of maximum proximity of model generated LI to empirical LI concomitant with matching of model estimated frequency and ASSR frequency gives us the confidence about the model estimation process when a complex range of frequencies are present in signal, e.g., during speech and melody processing. Furthermore, there could be different number of target regions in each frequency bands across different conditions. Hence, for practical reasons and to standardize the comparative analysis across conditions, empirical distribution of LI values across the entire brain volume (34 values) are correlated. Thus, the combination of *f*_*opt*_, *k*_*opt*_, *τ*_*opt*_ that yielded maximum correlation with empirical LI was selected for each participant (Fig. S1 B shows the stability and robustness if optimal parameters estimated beyond maximum correlation between empirical and simulated LIs). Note that the *k*_*opt*_ and *τ*_*opt*_ are valid only if the *f*_*opt*_ lies within the same range of empirical frequency band.

Furthermore, to confirm the accuracy of the model in utilizing structural connectivity (SC) to guide predictions, a control condition was implemented. In this condition, the weights and fiber lengths of the SC matrices for each participant were shuffled and used for the connectome model simulation. Specifically, the weights and fiber lengths of the SC matrices were randomly reassigned while maintaining the same number of connections and nodes. Comparative analyses were then performed between the shuffled and empirical 40 Hz ASSR condition, focusing on the assessment of the frequency specificity of the model. In particular, the model’s ability to accurately predict the frequency-specific responses, hence at 40 Hz, was examined. The presence of frequency-specific enhancement (*f*_*opt*_) in the shuffled and unshuffled conditions was compared, with particular emphasis on the distribution of *f*_*opt*_ in each condition. By contrasting the shuffled and unshuffled conditions, this assessment aimed to determine the extent to which the model’s predictions were reliant on the intact organization of the structural connections in the brain. Notably, the fact that each task state such as processing speech / melody requires idiosyncratic distribution of these free parameters given the specific nature of information processing. Additionally, these neurophysiologically relevant free parameters would not be identical in every participant owing to the inter-participant variability (Seghier & Price, 2018). participant-specific prediction enables not only contribute to the robustness of the analysis but also enables attributing the functional variability to the differences in individual structural connectivity. Nevertheless, another attribute of population studies is the existence of central tendency in the distribution of free parameter across participants. Hence, assessing the normality of *k*_*opt*_ and *τ*_*opt*_ across participants was required in this framework to evaluate the suitability of assuming a normal distribution for these parameters. A normal distribution indicates a predictable and stable pattern of parameter values, which is desirable for interpreting and generalizing the results of the model. Hence, the extent of prediction for individual regions were assessed and its inter-participant variability as the empirical source analysis revealed that only specific regions exhibited frequency-specific enhancement in Granger causality (GC) outflow from the bilateral PAC. Linear regression analysis was employed, wherein the empirical LI values of individual regions across participants were regressed against the corresponding simulated LI values. The goodness of fit was evaluated by assessing the coefficient of determination (R-squared) and the significance of the regression coefficients. The LI values of a particular region were selected from the spatial distribution of LI that yielded maximum correlation at *f*_*opt*_, *k*_*opt*_, and *τ*_*opt*_. Finally, the global coupling (*k*_*opt*_) and mean delay (*τ*_*opt*_) were examined between the speech and melody conditions across all frequencies via paired t-test. This analysis allowed to explore whether the brain exhibited distinct functional mechanisms to process auditory signals of various spatiotemporal complexities, e.g., speech, melody and rhythmic tonal sounds.

## Data and code availability

Anonymised and preprocessed data along with the analysis and simulation scripts are available at https://osf.io/6xk9f/.

## Acknowledgments

Authors acknowledge NBRC Core funds, Computing facility and Prof Robert Zatorre (Montreal Neurological Institute) for helpful suggestions. Authors also acknowledge the Neuroscience Gateway (Sivagnanam et al., 2013) for facilitating the resources required for model simulation.

## Funding

Ministry of Youth Affairs and Sports grant F.NO.K-15015/42/2018/SP-V(AB)

Department of Biotechnology Flagship program BT/MED-III/NBRC/Flagship/Flagship2019 (AB, DR)

## Author contributions

Conceptualization: NJ, AB

Methodology: NJ, PA, DR, AB

Investigation: NJ, AB.

Visualization: NJ

Supervision: DR, AB

Writing—original draft: NJ and AB

Writing—review & editing: NJ, PA, DR, AB

## Supplementary Materials

## SUPPLEMENTARY INFORMATION

**This supplementary analysis demonstrates the stability of the estimated global coupling (*k*_*opt*_) and global transmission delay (*τ*_*opt*_) parameters during distinct auditory conditions based on the model’s accuracy in capturing frequency-tuned neural dynamics (*f*_*opt*_).**

## Method

### Visualization of correlation gradient across parameter space

To depict the model’s ability to accurately capture frequency-specificity of neural dynamics and to visualize the landscape of model fit, we plotted values of the correlation coefficients between empirical and simulated Laterality Indices (LI) across a subset of parameter space of the Kuramoto oscillator network simulation. This visualization was done for the 40 Hz Auditory Steady-State Response (ASSR) condition, using data from a single participant.

### Validity of estimated parameters beyond maximum correlation coefficient

To further assess the robustness and stability of the estimated optimal parameters (*f*_*opt*_, *k*_*opt*_, *τ*_*opt*_), an analysis was conducted to examine their behavior across the top-ranked correlation coefficients, beyond the global maximum. Therefore, parameter combinations corresponding to the top 20 highest correlation coefficients from model fitting were identified for each subject. Thereafter, for each of the top 20 ranks, the Euclidean distance of the corresponding parameters from the top optimal parameter set *f*_*opt*_, *k*_*opt*_, *τ*_*opt*_ was computed separately for frequency (*f*), global coupling (*k*), and time delay (τ). Additionally, the corresponding correlation coefficients at each of these 20 ranks were tracked. This analysis aimed to demonstrate the stability of the identified optimal parameters even when selecting from suboptimal correlation coefficients, providing insight into the “flatness” or “sharpness” of the correlation landscape around the global optimum.

**Figure S1:**
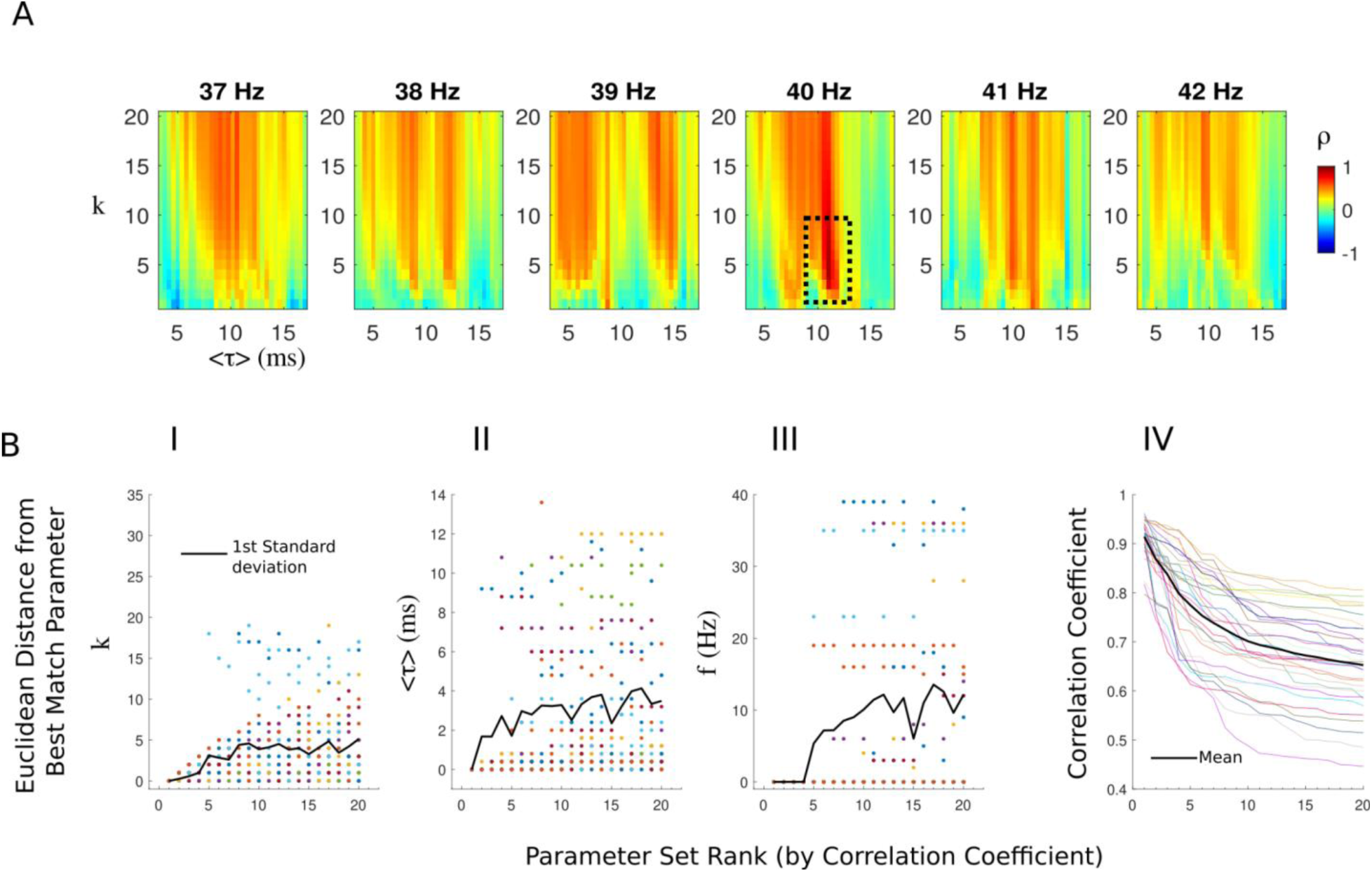
**A.)** Gradient of correlation coefficients across the entire parameter space of k, tau and subset of frequencies (37-42 Hz) from a single participant matched for 40 Hz ASSR condition. Each subplot represents input frequency of auditory nodes in the model simulation. The x-axis represents transmission delays (tau), and the y-axis represented global coupling (k), with the color intensity indicating the strength of correlation coefficient. **B.)** Panels I-III show the Euclidean distance of *k*, τ, and *f* (respectively) from their optimal values, plotted against the rank (1st to 20th). Each point represented a single participant, and a black line indicates the standard deviation across participants at each rank. Panel IV show the value of correlation coefficients at each of the top 20 ranks. Each color line represents a single participant, and a black line indicated the mean correlation coefficient across participants at each rank.

## Results

### Frequency-specific based optimal parameter identification

The visualization of the correlation coefficients across the parameter space for the representative participant clearly demonstrated a global maximum in model-to-empirical fit, shown as the dashed box in the Figure S1 A. Specifically, the maximum Correlation coefficient (dashed box) was present at transmission delays (τ*opt* = 11 ms), global coupling (*kopt* = 4) and frequency (*fopt* = 40 Hz). Since, the optimal frequency (*fopt*) derived from the model fitting process precisely matches the empirical 40 Hz frequency of the ASSR validates the identified optimal global coupling (*k*_*opt*_) and transmission delay (*τ*_*opt*_) parameters are biologically valid and mechanistically relevant. The present result provides strong validation for the model’s accuracy and its ability to capture frequency-specific neural dynamics

### Validity of estimated parameters beyond maximum correlation coefficient

The analysis of optimal parameters stability across the top 20 correlation ranks (Figure S1 B) provided insights into the robustness of the identified optimal parameters (*f*_*opt*_, *k*_*opt*_, *τ*_*opt*_). The Euclidean distance of *k*_*opt*_ and *τ*_*opt*_ (Panels I-II) from their respective top *params_opt_* remained relatively stable across the top 20 correlation ranks. The black line, representing the standard deviation among participants, showed a gradual increase, indicating that even when selecting slightly suboptimal correlation coefficients, the underlying k and tau values did not drastically deviate from their optimal counterparts given the massive parameter space. Though, the input frequency (*f*) showed a sudden increase in the Euclidean distance from its optimal value (*fopt*) after approximately the 4th top correlation coefficient. This indicates that while the very top correlation coefficients are tightly clustered around the optimal frequency, moving further down the rank list quickly leads to frequencies that are less specific to the 40 Hz ASSR. This sharp change highlights the range of frequency specificity of the model’s fit to the ASSR data. The plot of correlation coefficients at each rank showed a expected general decrease as the rank moved from 1st (maximum) to 20th.

